# Test-retest reliability of functional brain network characteristics using resting-state EEG and graph theory

**DOI:** 10.1101/385302

**Authors:** Bahar Moezzi, Brenton Hordacre, Carolyn Berryman, Michael C. Ridding, Mitchell R. Goldsworthy

## Abstract

Metrics of brain network organization can be derived from neuroimaging data using graph theory. We explored the test-retest reliability of graph metrics of functional networks derived from resting-state electroencephalogram (EEG) recordings. Data were collected within two designs: (1) within sessions (WS) design where EEG data were collected from 18 healthy participants in four trials within a few hours and (2) between sessions (BS) design where EEG data were collected from 19 healthy participants in three trials on three different days at least one week apart. Electrophysiological source activity was reconstructed and functional connectivity between pairs of sensors or brain regions was determined in different frequency bands. We generated undirected binary graphs and used intra-class correlation coefficient (ICC) to estimate reliability. We showed that reliabilities ranged from poor to good. Reliability at the sensor-level was significantly higher than source-level. The most reliable graph metric at the sensor-level was cost efficiency and at the source-level was global efficiency. At the sensor-level: WS reliability was significantly higher than BS reliability; high beta band in WS design had the highest reliability; in WS design reliability in gamma band was significantly lower than reliability in low and high beta bands. At the source-level: low beta band in BS design had the highest reliability; there was no significant main effect of frequency band on reliability; reliabilities of WS and BS designs were not significantly different. These results suggest that these graph metrics can provide reliable outcomes, depending on how the data were collected and analysed.

## 1. Introduction

The integration of information between functionally specialised and widely distributed brain areas is important for complex behaviour (Bressler and Menon 2010; Davis *et al*. 2012; Smith *et al*. 2015). Functional connectivity reflects the temporal correlations of neural activity between brain areas and can be estimated using EEG recordings (Horwitz 2003). Resting-state brain activity is suggested to exhibit valuable information on how various areas of the brain communicate (Deco *et al*. 2011). Graph Theory is an appropriate framework for the mathematical treatment of complex networks. The complex brain networks can be represented as graphs in which nodes correspond to the EEG electrodes or brain regions and edges to the functional connectivity between them (Bullmore and Sporns 2009; Rubinov and Sporns 2010). Graph metrics have been used to quantify the modification of the functional organization of the brain by age (Wang *et al*. 2010), sex (Tian *et al*. 2011), genetic disposition (Fornito *et al*. 2011), and mental disease (Rubinov *et al*. 2009; Lynall *et al*. 2010).

Before the utility of graph metrics of brain networks derived from EEG data can be fully appreciated as biomarkers of brain function, it is important to consider reliability of their repeated measurements in the same subjects at different frequency bands. Previous studies reported that the reliabilities of the graph metrics of resting-state functional connectivity derived from magnetoencephalography (MEG) data at the sensor-level and from functional magnetic resonance imaging (fMRI) data range from poor to excellent (Cao *et al*. 2014; Jin *et al*. 2011; Braun *et al*. 2012; Deuker *et al*. 2009). Hardmeier *et al*. (2014) recorded resting-state EEG data in three trials separated by one year and reported poor to excellent reliability for graph metrics: small world index, normalized clustering coefficient, normalized average path length and regional degree in different frequency bands. They computed these metrics at the sensor level based on the connectivity measures phase-lag index (PLI) and weighted PLI. Recently, Kuntzelman and Miskovic (2017) computed connectivity measures spectral coherence and debiased weighted PLI from resting-state EEG data recorded in two trials separated by one week. They computed graph metrics: global efficiency, characteristic path length, radius, diameter, modularity, transitivity, strength, local efficiency, clustering coefficient, betweenness centrality, eigenvector centrality and page-rank centrality at the sensor-level. They reported poor to excellent reliability for these metrics in different frequency bands. These studies suggest that some graph metrics derived from resting-state EEG recordings at the sensor-level can be utilized in a reliable manner in basic and clinical research settings. However, the interpretation of connectivity measures from sensor-level EEG data is not straightforward. Instead, the electrophysiological sources of sensor-level data can be reconstructed, facilitating a source-level approach in graph theoretical analysis of EEG data (Babiloni *et al*. 2004).

In this study, we investigated test-retest reliability of graph metrics estimated from resting-state EEG data at both sensor- and source-levels in both short-term (within a few hours, i.e. WS) and long-term (several days apart, i.e. BS) designs at different frequency bands. Imaginary coherence was used to construct functional networks because it effectively suppresses spurious coherence driven by field spread (Nolte *et al*. 2008). Undirected binary graphs were used to compute graph metrics: betweenness centrality, cost efficiency, characteristic path length, global efficiency, local efficiency, clustering coefficient, k- coreness, and maximized modularity. Test-retest reliability of these graph metrics and also mean imaginary coherence was quantified over all pairs of electrodes (sensor-level) or brain regions (source-level) using intra-class correlation coefficient (ICC).

This study is significant for the field of neuroimaging as it will help establish the reliability of EEG-based graph metrics in order to incorporate them into future designs and facilitate appropriate selection of reliable estimates of functional connectivity.

## 2. Materials and Methods

### 2.1. Participants

We used data collected from a total of 23 healthy participants (9 male, 22 right handed, mean aged 25 (SD 7) years). EEG was recorded from 18 participants for WS design and from 19 participants for BS design, with 14 participants attending both designs. All participants gave written informed consent in accordance with the World Medical Association Declaration of Helsinki to participate in this study. Ethical approval was provided by the University of Adelaide Human Research Ethics Committee.

### 2.2. Experimental Design

For this study, we analysed resting-state EEG data (eyes-open condition). Participants were instructed to view a fixation point, remain as still, quiet and relaxed as possible and if possible, try and avoid blinking too much. The EEG recordings were part of measurements embedded in two studies: a study on effects of continuous theta burst stimulation (cTBS) on the brain dynamics (WS and BS) and a study on effects of pain induction on the brain dynamics (BS only). Only pre-intervention or sham condition data were used for these analyses. In the WS design, for each participant, data were collected in four trials in one day on average 24 (SD 12) minutes apart as a part of a sham cTBS protocol, which does not apply current to the scalp or alter neural activity. In the BS design, data were collected from each participant at three separate experimental sessions separated by a minimum of 7 days (average 31, SD 41 days).

### 2.3. EEG acquisition and pre-processing

We acquired three minutes of continuous resting-state EEG using an ASA-lab EEG system (ANT Neuro, Enschede, Netherlands) or a TMSi EEG system (Twente Medical Systems International B.V, Oldenzaal, The Netherlands) using a WaveguardTM original cap with 64 sintered Ag-AgCl electrodes in standard 10-10 positions. Each individual’s test-retest session was performed on the same EEG system. Signals were sampled at 2048 Hz, amplified 20 times, filtered (high pass, DC; low pass 553 Hz) and referenced to the average of all electrodes. Impedance was kept below 5 k and the recorded data were stored on a computer for online analysis. EEG data were exported to MATLAB 9.0 (MathWorks, Inc., Natick, MA) for pre-processing and analysis. The signal was segmented into epochs of 1 s and channel baseline means were removed from the EEG dataset. Channels that were disconnected during recording or dominated by exogenous artefact noise were removed, and data were filtered using a hamming windowed sinc FIR (Finite Impulse Response) filter (1-45 Hz). We detected and excluded epochs contaminated by excessive noise by a procedure based on the identification of theshold for the maximum allowed amplitude for the EEG signals (>80 μV, total rejection rate was 18%). Fast ICA artefact correction was used in order to correct for non-physiological artefacts (e.g. eye blinks and scalp muscle activity) (Delorme and Makeig 2004). Following artefact removal, missing channels were interpolated using super-fast spherical interpolation.

### 2.4. Source reconstruction

To develop a source reconstruction model a forward model was constructed (see Figure 1). The first step in constructing the forward model is to find the brain surface from the subject’s MRI. In the absence of individual MRI, detailed anatomical information was incorporated in the form of an MRI template. We used ICBM152 template ((Fonov *et al*. 2011), a non-linear average of the MRI images of 152 individual heads) in Montreal Neurological Institute (MNI) coordinate. Next, a segmentation procedure was run in which each of the voxels of the anatomical MRI was assigned to a tissue class (returning probabilistic gray matter/white matter/cerebrospinal fluid (CSF) masks). A volume conduction model specifies how currents that are generated by sources in the brain are propagated through the tissues and how these result in externally measurable EEG potentials. A volume conduction model was constructed from the geometry of the head using boundary element method (BEM, (Geselowitz 1967; Fuchs *et al*. 2001)) with OpenMEEG software ((Gramfort *et al*. 2010), freely available at http://openmeeg.github.io/). We used a standard EEG electrode placement that matched our EEG configuration ((Oostenveld *et al*. 2011), freely available at http://www.fieldtriptoolbox.org/template/electrode) and confirmed that EEG electrodes were aligned with the head model. Based on the segmented MRI and restricted to gray matter, the brain was discretised into a source grid of 3990 voxels. For each grid point the lead field matrix, expressing the forward solution, was calculated and normalized in order to control against the power bias towards the centre of the head. The spatially adaptive filters for each grid point in each frequency band were computed using partial canonical correlation (PCC) method ((Schoffelen *et al*. 2008), regularization parameter λ = 5%, fixed orientation). This was computed based on the cross spectral density matrices (fast Fourier transform multitaper approach, smoothing ± 2 Hz, discrete prolate spheroidal sequences as tapers). For our source analysis, we used FieldTrip which is a MATLAB software toolbox for EEG and MEG analysis (Oostenveld *et al*. 2011).

**Figure 1:**
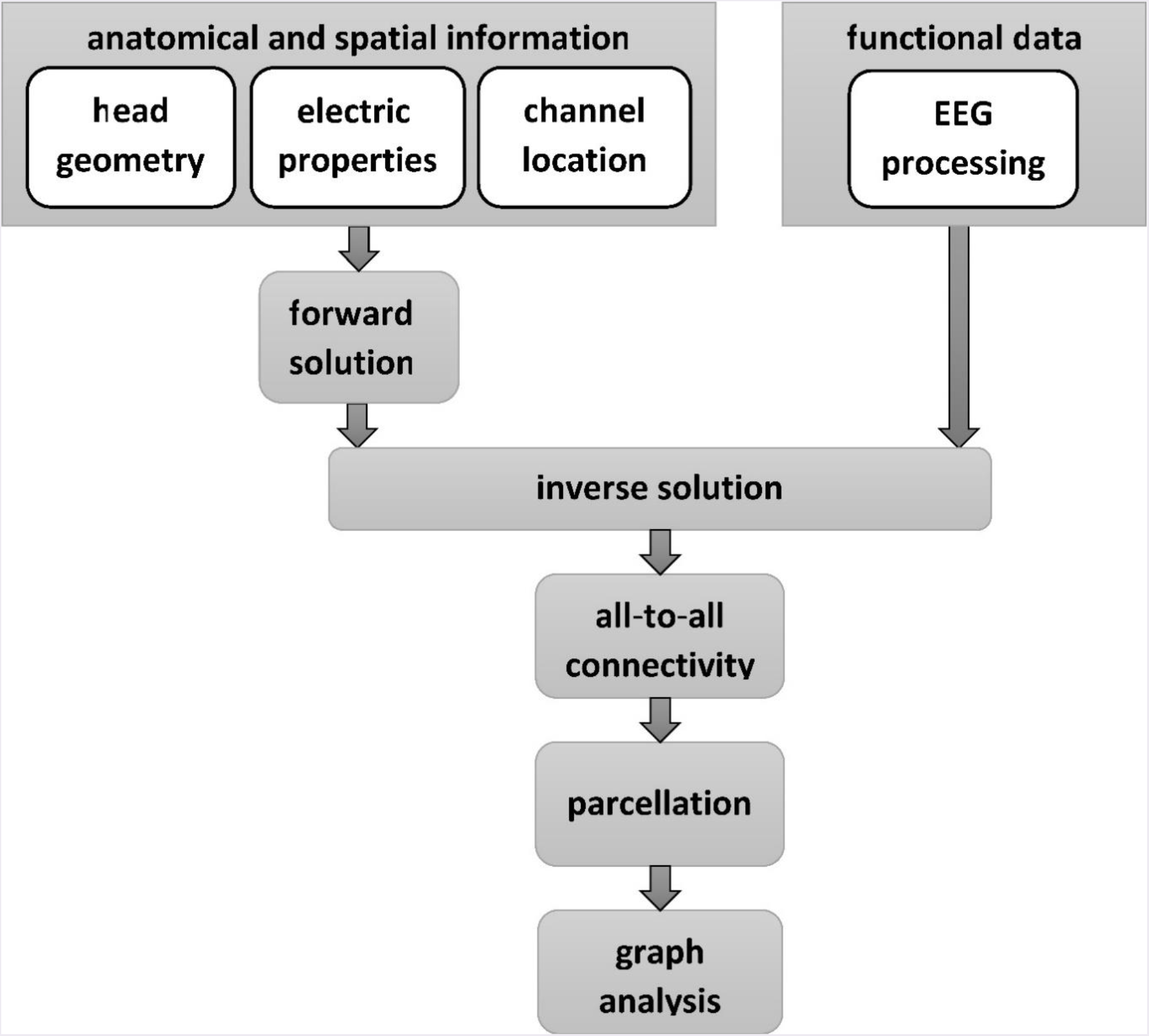
Schematic of the quantitative analysis.

### 2.5. Connectivity

We constructed the functional connectivity matrices using imaginary coherence. Denote the *l*-th segment of the *i*-th time course by *X_i;l_* and its Fourier transform by *X_i;l_*. The output of the PCC implementation contains the single trial estimates of amplitude and phase (Fourier coefficients). The cross spectral matrix is defined as

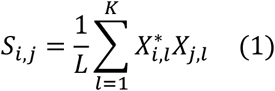

where (.)* denotes complex conjugation and *L* denotes the number of segments. Coherency between the *i*then co-registered to FieldT rip template then co-registered to FieldT rip template then co-registered to FieldT rip template then co-registered to FieldT rip template then co-registered to FieldT rip template -th and *j*-th times series is defined as

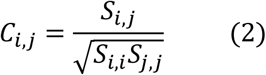

In order to define ROIs for source-level analysis, we first tesselated the cortical mesh within Brainstorm application (Tadel *et al*. 2011) using the Desikan-Killiany atlas (Desikan *et al*. 2006). This mesh was then co-registered to FieldTrip template mesh using the minimum Euclidean distance approach. We computed the absolute value of imaginary coherence between two ROIs *p* and *q* as

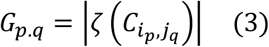

where 𝜁(.) denotes the imaginary part and *i*_p_-th and *j*_q_-th voxels are at the geometric centres of p and q, respectively. Centre voxels refer to those voxels that have minimum Euclidean distance with all other voxels within the ROI. Finally, matrix G is divided by its standard deviation using Jackknife method to produce a connectivity matrix. Similarly, at the sensor-level, the absolute value of imaginary coherence between each two sensors was computed.

### 2.6. Graph theory analysis

We thresholded the connectivity matrices and constructed binary undirected networks. The connectivity matrices were tested over a range of different thresholds to cancel the issue of semi-arbitrary thresholding (Langer *et al*., 2013). Network density is defined as the percentage of the connections (edges) that remain after thresholding. The network density was changed to control the threshold levels. The connection density of 10% was tested as this was reported to provide an optimal trade-off between reducing spurious connections and retaining true connections providing biologically plausible information about the brain functional networks (Dosenbach *et al*. 2010; Lord *et al*. 2012; Pedersen *et al*. 2015; Cocchi *et al*. 2015). For the graph theory analysis, we used Brain Connectivity Toolbox (BCT, (Rubinov and Sporns 2010), freely available at https://sites.google.com/site/bctnet/Home). The following graph metrics were computed for each subject for frequency bands theta (4-7 Hz), alpha (8-13 Hz), low beta (14-20 Hz), high beta (21-30 Hz) and gamma (31-45 Hz). We subsequently explored the effects of density on reliability by estimating all of these graph metrics at connection densities of 5% and 15%.

### 2.6.1. Global efficiency

Global efficiency is defined as the average inverse shortest path length and can be regarded as a measure of global integration (Latora and Marchiori 2001).

### 2.6.2. Local efficiency

Local efficiency is defined as the efficiency of the local sub-graph of a given node that contains only the direct neighbours of the node. It is a measure of local connectedness (Latora and Marchiori 2001).

### 2.6.3. Betweenness centrality

Betweenness centrality is defined as the number of shortest paths that run through a given node. When averaged over all nodes, it is a measure of the influence of a node over information ow among other nodes in the network (Freeman 1977).

### 2.6.4. Cost efficiency

Cost efficiency is evaluated as

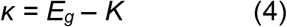

where E_g_ denotes global efficiency and *K* denotes network density (Achard and Bullmore 2007).

### 2.6.5. k-coreness

The k-core is defined as the largest subgraph comprising nodes of degree at least k. The coreness of a node is k if the node belongs to the k-core but not to the (k+1)-core (Hagmann *et al*. 2008).

### 2.6.6. Maximized modularity

The optimal community structure is a sub-division of the network into non-overlapping groups of nodes in a way that maximizes the number of within-groups edges and minimizes the number of between-groups edges. Modularity is a statistic that quantifies the degree to which the network may be sub-divided into such structure (Newman 2006).

### 2.6.7. Characteristic path length

Characteristic path length is defined as the average shortest path length in the network. The shortest path length between two nodes is the minimum number of edges between two nodes (Sporns *et al*. 2004).

### 2.6.8. Clustering coefficient

Clustering coefficient is defined as the proportion of the nearest neighbours of a given node that are connected to each other. It is a measure of local connectedness (Rubinov and Sporns 2010; Watts and Strogatz 1998).

### 2.7. Statistical analysis

The graph metrics were log-transformed prior to analyses to achieve normality. For each graph metric derived from the first EEG trial of the WS design, we tested for statistical differences between frequency bands at density 10% using repeated ANOVA.

ICC (two-way mixed single measures: testing for consistency) was estimated as a measure of test-retest reliability between the trials for each design (BS or WS) and each graph metric at each frequency band and density at both sensor and source-levels as defined in (Shrout and Fleiss 1979). Additionally, the ICCs were estimated based on the average of the elements of the connectivity matrix (mean imaginary coherence) independently of the density level. The negative ICCs were set to zero as suggested in other test retest studies using ICC (Braun *et al*. 2012; Kong *et al*. 2007). Each ICC was categorized into a value of >0.75 as “excellent”, 0.60-0.74 as “good”, 0.40-0.59 as “fair” and <0.40 as “poor” reliability (Cicchetti 1994).

ICCs were reported descriptively also using ANOVA models to test the effects of different factors on test-retest reliability of the metrics. Where Mauchly’s test indicated that the assumption of sphericity was violated, Greenhouse-Geisser tests were reported. All statistical analyses were performed using IBM SPSS Statistics 24 and MATLAB 9.0. Significance was measured at p<0.05.

## 3. Results

### 3.1. Sensor-level

#### 3.1.1. Comparison of the metrics between frequency bands

There was a significant main effect of frequency band on cost efficiency, global and local efficiency, clustering coefficient, k-coreness and mean imaginary coherence (see Table 1). Bonferroni post hoc tests revealed that: cost efficiency in theta band was significantly lower than cost efficiency in the alpha (p=0.04) and gamma (p=0.02) bands; global efficiency in theta band was significantly lower than global efficiency in the alpha (p=0.01) and gamma (p=0.01) bands; global efficiency in low beta band was significantly lower than global efficiency in alpha band (p=0.01); local efficiency in theta band was significantly lower than local efficiency in the high beta (p=0.02) and gamma (p=0.03) bands; clustering coefficient in theta band was significantly lower than in the high beta (p=0.01) and gamma (p=0.01) bands. k-coreness in alpha band was significantly higher than k-coreness in the theta (p=0.04) and gamma (p=0.02) bands; mean imaginary coherence in alpha band was significantly higher than mean imaginary coherence in the theta (p<0.001), low beta (p<0.001), high beta (p<0.001) and gamma (p<0.001) bands. There was no significant main effect of frequency band on betweenness centrality, characteristic path length and maximized modularity.

**Table 1.** Main effect of frequency band on different metrics. Bold font indicates significance (p<0.05).

|  | sensor-level<br>F-ratio | sensor-level<br>p-value | source-level<br>F-ratio | Source-level<br>p-value |
| --- | --- | --- | --- | --- |
| Betweenness centrality | 1.02 | 0.38 | 0.91 | 0.47 |
| Cost efficiency | **3.41** | 0.01 | 2.56 | 0.07 |
| Characteristic path length | 2.46 | 0.06 | 0.86 | 0.49 |
| Global efficiency | **5.77** | 0.004 | 2.52 | 0.07 |
| Local efficiency | **4.58** | 0.01 | 1.29 | 0.28 |
| Clustering coefficient | **5.41** | 0.01 | 0.97 | 0.43 |
| K-coreness | **10.61** | 0.01 | 0.62 | 0.65 |
| Maximized modularity | 1.07 | 0.37 | 1.28 | 0.33 |
| Mean imaginary coherence | **32.31** | <0.001 | **11.27** | <0.001 |

#### 3.1.2. Range of reliabilities

The ICC values for the graph metrics range from 0 to 0.48 (mean 0.25, SD 0.12) for WS design, and range from 0 to 0.41 (mean 0.13, SD 0.12) for BS design, indicating poor to fair reliability. The ICC values for mean imaginary coherence range from 0.17 to 0.71 (mean 0.42, SD 0.22) for WS design, and range from 0.30 to 0.49 (mean 0.41, SD 0.07) for BS design, indicating poor to good reliability (see Figure 2).

**Figure 2:**
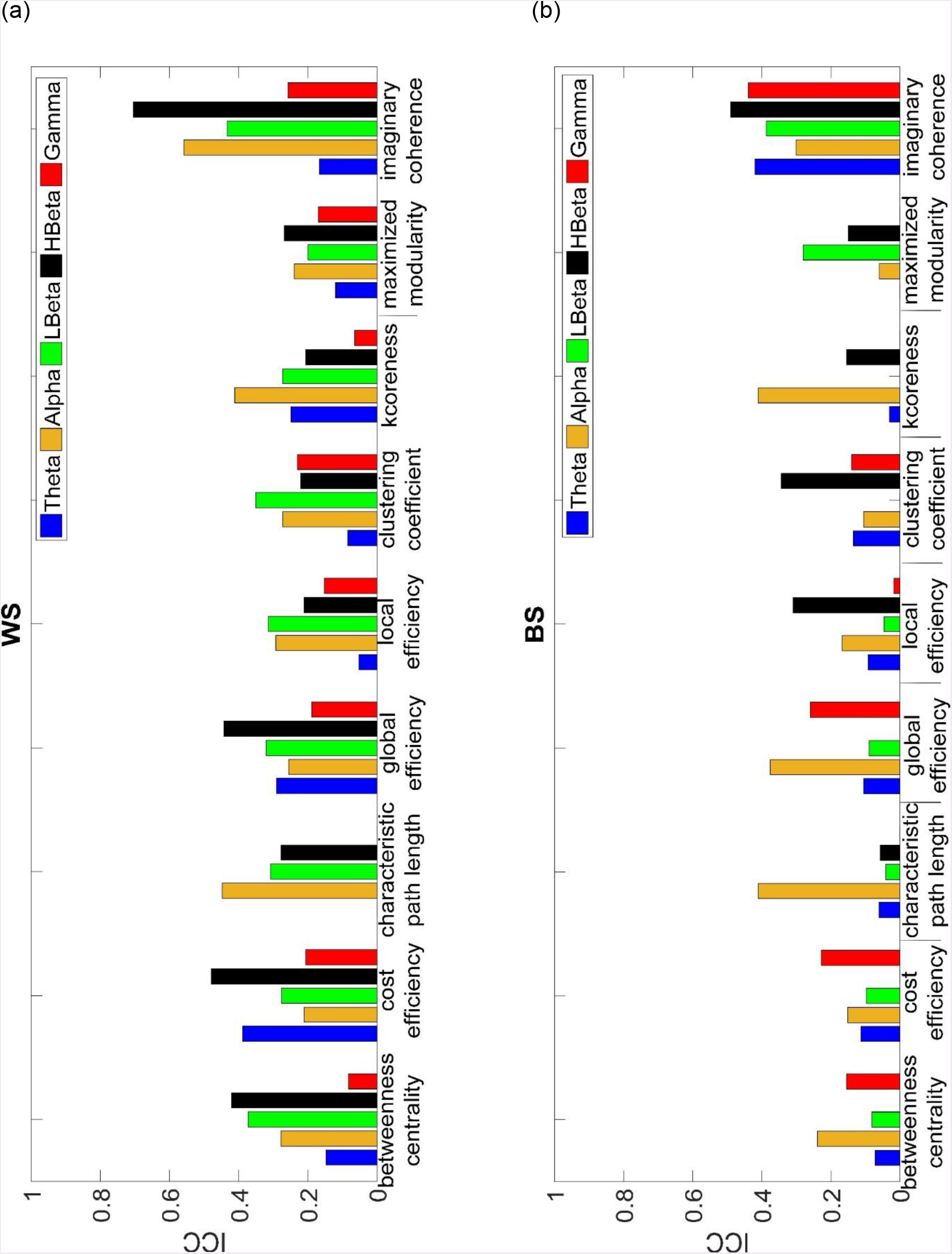
Test-retest reliability (ICC) of different metrics in (a) WS and (b) BS designs at the sensor-level in theta, alpha, low beta, high beta and gamma frequency bands.

#### 3.1.3. Comparison of reliabilities between frequency bands

Reliability varied over different frequency bands. The lowest mean ICC over all metrics in each of WS and BS designs in each frequency band was found in the low beta frequency band of the BS design (0.12±0.13(SD)), while the highest mean ICC was found in the high beta frequency band in the WS design (0.36±0.17(SD)) followed by alpha band in the WS design (0.33±0.12(SD)). Accordingly, an ANOVA model that treated the different frequency bands as within-subjects factor was used. There was a significant main effect of frequency band in WS design (F(4,32)=8.24,p=0.01). Bonferroni post hoc tests revealed that in WS design mean ICC in gamma band was significantly lower than mean ICC in the low beta band (p=0.01) and high beta band (p=0.03). No significant main effect of frequency band was observed for BS design (F(4,32)=1.89,p=0.14).

#### 3.1.4. Comparison of reliabilities between different metrics

Reliability also varied between different metrics. The lowest mean ICC over all frequency bands in each of WS and BS designs was found for maximized modularity in the WS design (0.20±0.07(SD)), indicating poor reliability and the highest mean ICC was found for mean imaginary coherence in the WS design (0.4±0.22(SD)) indicating fair reliability. The most reliable graph metric was found to be cost efficiency in WS design (0.26±0.17(SD)). The mean ICC over all frequency bands and designs for graph metrics (0.15±0.14(SD)) was lower than for mean imaginary coherence (0.35±0.21(SD)).

#### 3.1.5. BS versus WS reliability

Additionally, we compared the reliability of the metrics for WS design versus BS design. The mean ICC over all frequency bands and metrics of the WS design (0.27±0.14(SD)) was higher than BS design (0.16±0.15(SD)). A two factor repeated measures ANOVA with frequency band as a within-subjects factor and WS versus BS design as a group factor showed that WS reliabilities were significantly higher than the BS reliabilities (F(1,16)=7.28,p=0.02); excluding the ICCs of mean imaginary coherence (F(1,14)=49.27, p<0.001).

#### 3.1.6. Effects of network density on reliability

All the graph metrics were initially estimated in networks of 10% density. We subsequently explored the effects of density on reliability by estimating all metrics in networks with densities 5% and 15% (see Figure 3). There was a significant main effect of density in WS design (F(2,14)=7.12,p=0.01). Bonferroni post hoc tests revealed that reliability at 15% density was significantly lower than at 10% (p=0.048). In the BS design, there was no significant main effect of connection density (F(2,14)=1.79,p=0.23).

**Figure 3:**
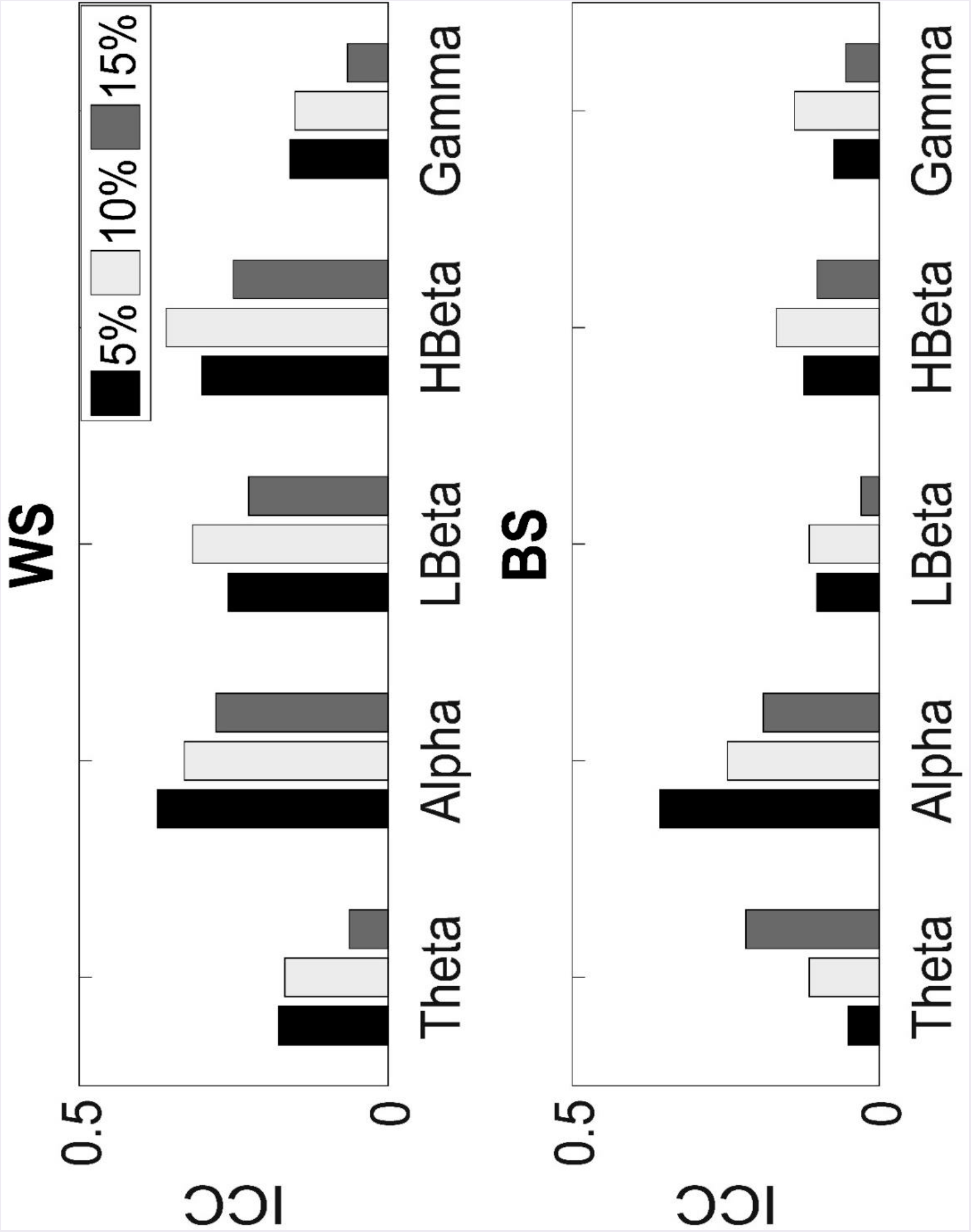
Mean test-retest reliability (ICC) over all the metrics at densities 5%, 10% and 15% in different frequency bands at the sensor-level in the WS and BS designs.

### 3.2. Source-level

#### 3.2.1. Comparison of the metrics between frequency bands

There was no main effect of frequency band on the graph metrics (See Table 1). However, there was a significant main effect of frequency band on mean imaginary coherence. Bonferroni post hoc tests revealed that the mean imaginary coherence in alpha band was significantly higher than mean imaginary coherence in the theta (p=0.02), low beta (p=0.01) and gamma (p < 001) bands and mean imaginary coherence in the gamma band was significantly lower than in the high beta band (p=0.02).

#### 3.2.2. Range of reliabilities

The ICC values for the graph metrics range from 0 to 0.26 (mean 0.03, SD 0.06) for WS design and range from 0 to 0.28 (mean 0.07, SD 0.07) for BS design, indicating poor reliability. The ICC values for mean imaginary coherence range from 0.23 to 0.49 (mean 0.31, SD 0.10) for WS design and range from 0.10 to 0.30 (mean 0.23, SD 0.81) for BS design indicating poor to fair reliability (Figure 4).

**Figure 4:**
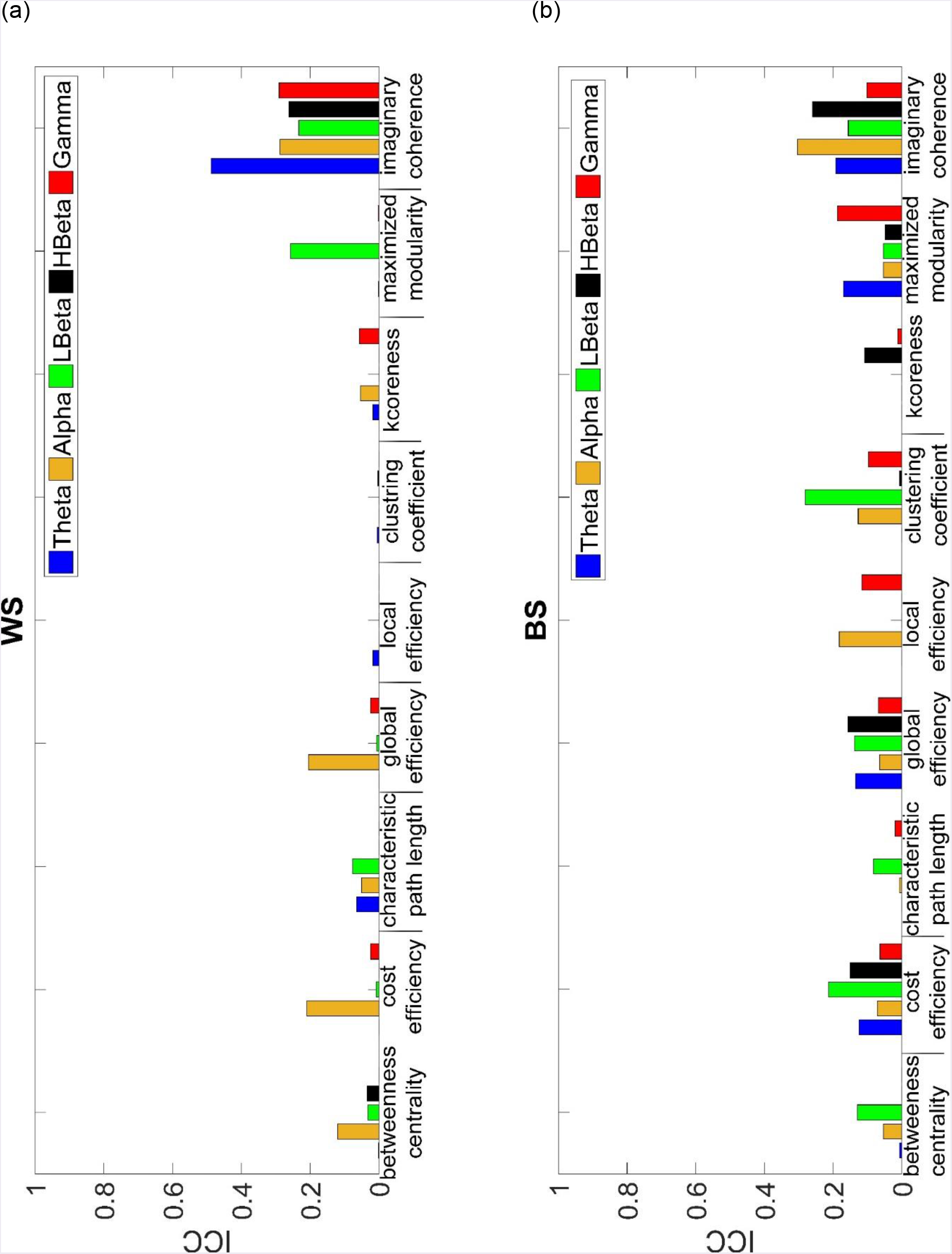
Test-retest reliability (ICC) of different metrics in (a) WS and (b) BS designs at the source-level in theta, alpha, low beta, high beta and gamma frequency bands.

#### 3.2.3. Comparison of reliabilities between frequency bands

Reliability varied over different frequency bands. The lowest mean ICC over all metrics in each of WS and BS designs in each frequency band was found in the high beta frequency band of the WS design (0.03±0.08(SD)). The highest mean ICC was found in the low beta frequency band in the BS design (0.12±0.09(SD)) followed by alpha band in WS design (0.05±0.04(SD)). Accordingly, an ANOVA model that treated the different frequency bands as within-subjects factor was used. No significant main effect of frequency band was observed for WS design (F(2.31,18.48)=1.38,p=0.28). Similarly, no significant main effect of frequency band was observed for BS design (F(4,32)=0.66,p=0.63).

#### 3.2.4. Comparison of reliabilities between different metrics

Reliability also varied between different metrics. The lowest mean ICC over all frequencies in each of WS and BS designs was found for clustering coefficient in the WS design (0.002±0.002(SD)) and the highest mean ICC was found for mean imaginary coherence in the WS design (0.31±0.10(SD)). The most reliable graph metric was found to be global efficiency in BS design (0.04±0.08(SD)). The mean ICC over all frequency bands and designs for graph metrics (0.04±0.07(SD)) was lower than for mean imaginary coherence (0.22±0.14(SD)).

#### 3.2.5. BS versus WS reliability

Additionally, the reliability of the metrics was compared for WS design versus BS design. The mean ICC over all frequency bands and metrics in the BS design (0.09±0.08(SD)) was higher than WS design (0.06±0.11(SD)); however, a two factor repeated measures ANOVA with frequency band as a within-subjects factor and WS versus BS design as a group factor showed that the reliabilities of WS design were not significantly different from the BS reliabilities (F(1,16)=0.44,p=0.516); excluding the ICCs of mean imaginary coherence (F(2.54,35.55)=2.02,p=0.14).

#### 3.2.6. Effects of network density on reliability

Again, the effects of density on reliability were explored (see Figure 5). No significant main effect of connection density for WS design was observed (F(1.16,8.13)=2.73,p=0.135). For BS design, there was a significant main effect of density (F(2,14)=7.26,p=0.01). Bonferroni post hoc tests revealed that reliability at 15% density was significantly lower than at 10% (p=0.04).

**Figure 5:**
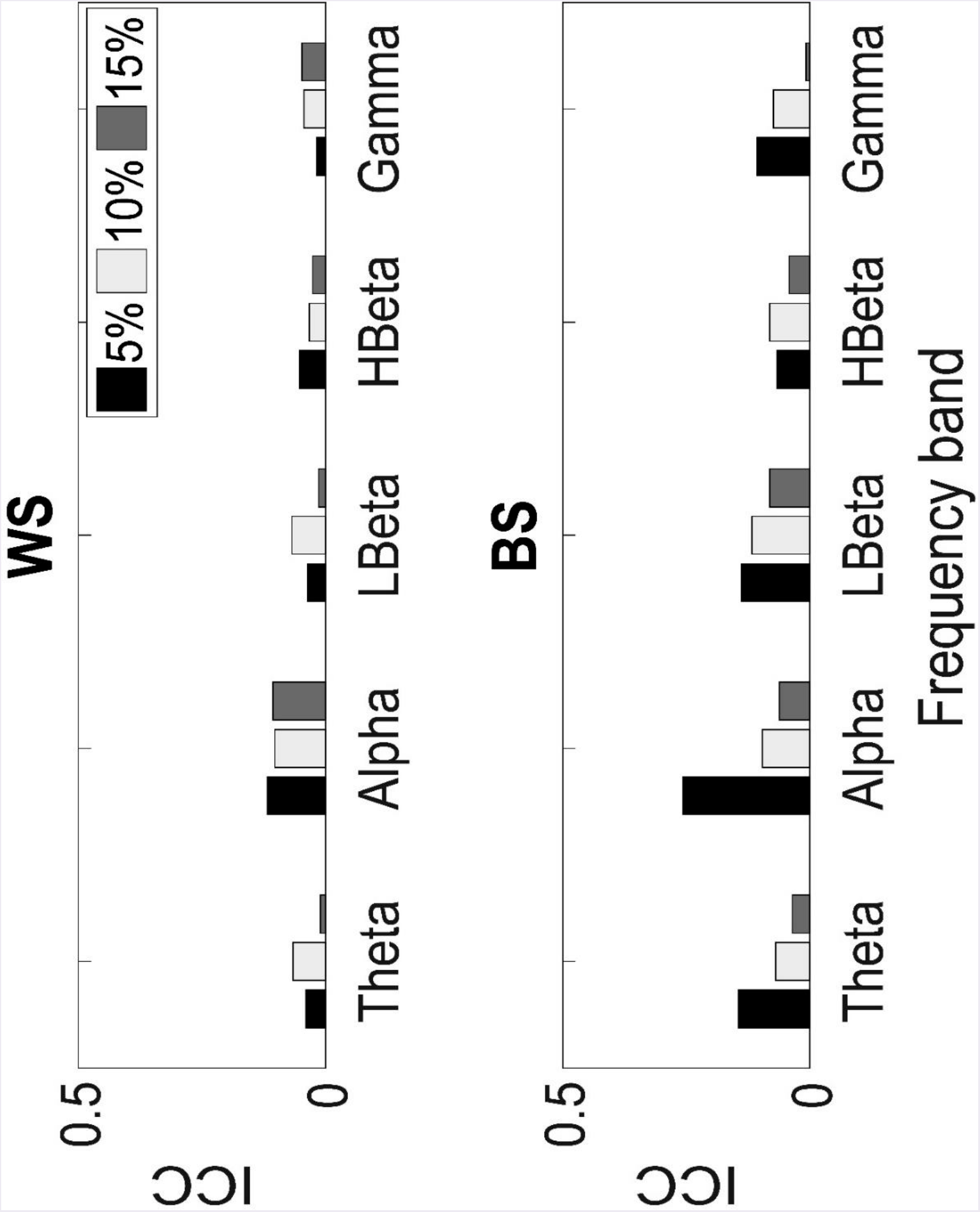
Mean test-retest reliability (ICC) over all the metrics at densities 5%, 10% and 15% in different frequency bands at the source-level in WS and BS designs.

### 3.3. Comparison of sensor-level and source-level reliabilities

The reliability of the sensor-level versus source-level metrics was compared at 10% density (see Figure 6). The mean ICC over all frequency bands, metrics and designs at the sensor-level (0.21±0.15(SD)) was higher than source-level (0.08±0.10(SD)). A two factor repeated measures ANOVA with frequency band and sensor/source-level as within-subjects factors and WS versus BS design as a group factor showed that sensor-level reliabilities were significantly higher than the source-level reliabilities (F(1,16)=101.56, p<0.001).

**Figure 6:**
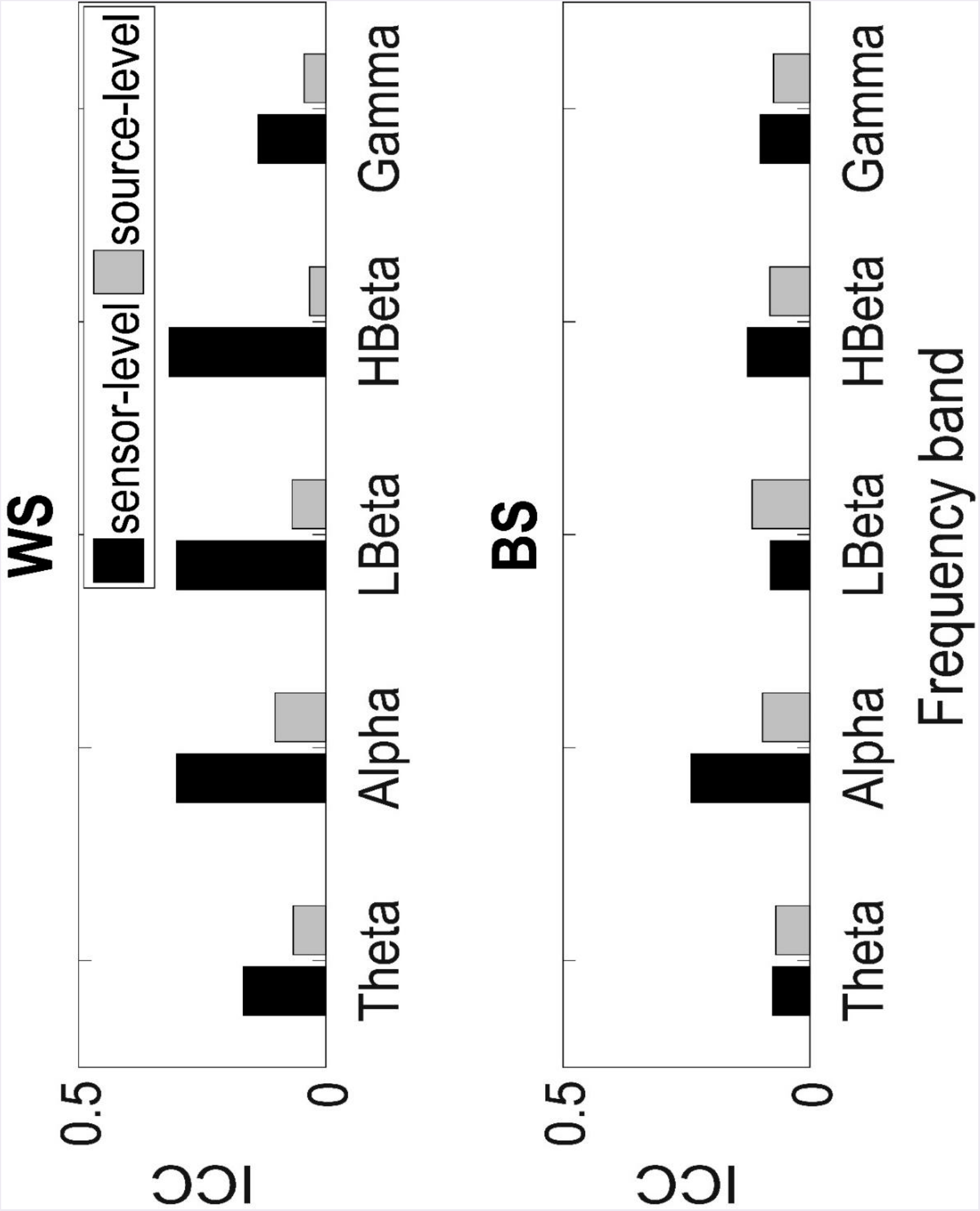
Mean test-retest reliability (ICC) over all the metrics at 10% density in different frequency bands at the sensor- and source-levels for WS and BS designs.

## 4. Discussion

The main purpose of our study was to quantify the reliability of sensor- and source-space graph metrics derived from resting-state EEG. We were particularly interested in comparing reliability of the graph metrics derived from data collected within a few hours with those derived from data collected at least a week apart. Using EEG data enabled us to also compare reliabilities at different frequency bands. Overall, we observed poor to good reliabilities for the mean imaginary coherence and eight graph metrics. For sensor-level outcomes, we observed levels of reliability that were comparable to those derived from resting-state EEG and MEG at the sensor-level in previous studies (Jin *et al*. 2011; Deuker *et al*. 2009; Hardmeier *et al*. 2014; Kuntzelman and Miskovic 2017). To our knowledge, the present study is the first analysis of test-retest reliability of graph metrics at both sensor- and source-levels derived from EEG data with both WS and BS designs.

The graph metrics varied across different frequency bands. This may suggest frequency-specific network organization and imply various functional roles for different frequency bands. The highest reliability averaged over all metrics at the sensor-level was found in the high beta frequency followed by alpha frequency band in the WS design. At the source-level the highest reliability averaged over all metrics was found in the low beta band in the BS design followed by alpha band in the WS design. This is in agreement with (Jin *et al*. 2011) which suggested that alpha and beta frequency bands provide the most reliable graph metrics in the resting-state. At the sensor-level in the WS design, reliability in the gamma band was significantly lower than low and high beta bands. This is consistent with findings in (Jin *et al*. 2011; Deuker *et al*. 2009) that the resting-state graph metrics are least reliable in the gamma band.

Characteristic path length (at the source-level) was found to be the least reliable graph metric, which is consistent with previous reports (Cao *et al*. 2014). Consistent with (Braun *et al*. 2012), the most reliable graph metric was found to be cost efficiency measured at the sensor-level. This potentially shows that this metric quantifies aspects of brain function that can be more reliably estimated using resting-state EEG.

It is expected that the greater the time between EEG data collection sessions, the larger the possibility of change in functional network organization of the brain and hence the lower the reliability of graph metrics. Accordingly, we found that the WS reliabilities were significantly higher than the BS reliabilities at the sensor-level, but interestingly, not significantly different at the source-level. We also showed that our sensor level reliabilities were significantly higher than the source-level reliabilities. An important concern when estimating functional brain network metrics from EEG recordings is the mixing of brain sources resulting in spurious results. Source mixing occurs at the sensor-level to a higher extent than at the source-level and is typically quite stable across participants and repeated measurements. As suggested by (Colclough *et al*. 2016), this is potentially one of the reasons sensor-level reliabilities were higher than the source-level reliabilities.

A potential contributor to the low reliability of some metrics in our study might be the relatively short EEG recording duration. Three-minute recording, as in the present study, might not be enough to provide reliable graph metrics. However, using four minutes of data, it was suggested that in general, duration of resting-state EEG makes little difference on reliability of the graph metrics (Kuntzelman and Miskovic 2017). Additionally, there are potential problems with recording EEG for long durations. During resting-state participants are required to maintain a constant state of arousal while refraining from moving. However, maintaining this state becomes more difficult with increased recording duration.

A limitation of our study is that our source-level results are specific to the methods we used for source reconstruction. We only looked at one (common) source reconstruction method. There is a large list of combinations of patient and EEG-setup, functional connectivity measures and network metrics and this study adds information on a portion. Mahjoory *et al*. (2017) reported a high level of variability in the functional connectivity estimates derived from common source reconstruction methods. Future work should compare the reliabilities of graph metrics obtained using different source reconstruction methods. A potential strength of the present study compared to previous literature is the relatively high number of trials, both in the BS and WS designs. However, the limitations include the relatively low density of EEG sensors compared to other similar studies (Deuker *et al*. 2009; Kuntzelman and Miskovic 2017) and low sample size. Our sample size and sensor density are commonly implemented in various research settings, and therefore are particularly relevant. Nevertheless, increasing the number of participants and utilizing higher density of EEG electrodes may yield better reliability, and this needs to be examined in future work. Another limitation of our study is that we used binary rather than weighted networks which might result in ignoring important information (Rubinov and Sporns 2011) and affect our results. Different EEG systems may have also influenced the outcome. However, different research laboratories use various systems; hence our results will be of interest to the research community. We reconstructed over 3000 sources from 64 electrodes. Future studies should consider changing the number of sources in order to cancel the effects of local under- or over-fitting. The use of a sham condition, which is considered to be a resting-state no-task condition in this study, may also have affected the outcome measures through placebo effects. Finally, the data in this study was not collected specifically for the intended analysis. Even though the data were collected in pre-intervention or sham intervention conditions in the original studies it is possible that carry over effects from the real (cTBS or pain) arms of the studies influenced the measures reported here. However, this is unlikely given that there was at least a seven-day period between conditions and there is no evidence to suggest any induced changes can last for so long (Huang *et al*. 2005; Nyfleler *et al*. 2006, 2009; Schabrun *et al*. 2015; Torta *et al*. 2017). However, it is important to repeat this study on datasets designed specifically for the intended analysis to confirm that reliability estimates were not affected by these experimental factors.

In conclusion, the connectivity and graph metrics derived from resting-state EEG data can provide robust results, although this differs depending on the frequency band, graph metric of interest, choice of sensor-level or source-level, and whether measurements are made within or between sessions. Sensor-level reliabilities were found to be higher than source-level reliabilities, particularly in the within-session design. Our source-level results highlight the need to take caution when interpreting source-level outcomes. Further work is needed to determine methodological approaches that could improve reliability of source measures, and to identify the factors responsible for intra-individual differences in EEG graph metrics.

## 5. Acknowledgement

MRG is supported by an NHMRC-ARC Dementia Research Development Fellowship (1102272). BH is supported by an NHMRC Early Career Researcher Fellowship (1125054). CB is supported by an NHMRC Early Career Researcher Fellowship (1127155). We would like to thank Stuart Howell of the University of Adelaide for statistical analysis support. BM would like to thank Luke Hallam for helpful discussions.

## References

Achard, S, Bullmore, E (2007) Effciency and cost of economical brain functional networks. PLoS Comput Biol 3: 174–183. doi:10.1371/journal.pcbi.0030017.

Babiloni, C, Binetti, G, Cassetta, E, Cerboneschi, D, Dal Forno, G, Del Percio, C, Ferreri, F, Ferri, R, Lanuzza, B, Miniussi, C, Moretti, DV, Nobili, F, Pascual-Marqui, RD, Rodriguez, G, Romani, GL, Salinari, S, Tecchio, F, Vitali, P, Zanetti, O, Zappasodi, F, Rossini, PM (2004) Mapping distributed sources of cortical rhythms in mild Alzheimer’s disease: A multicentric EEG study. Neuroimage 22: 57–67. doi:10.1016/j.neuroimage.2003.09.028.

Braun, U, Plichta, MM, Esslinger, C, Sauer, C, Haddad, L, Grimm, O, Mier, D, Mohnke, S, Heinz, A, Erk, S, Walter, H, Seiferth, N, Kirsch, P, Meyer-Lindenberg, A (2012) Test-retest reliability of resting-state connectivity network characteristics using fMRI and graph theoretical measures. Neuroimage 59: 1404–1412. doi:10.1016/j.neuroimage.2011.08.044.

Bressler, SL, Menon, V (2010) Large-scale brain networks in cognition: Emerging methods and principles. Trends Cogn Sci 14: 277–290. doi:10.1016/j.tics.2010.04.004.

Bullmore, ET, Sporns, O (2009) Complex brain networks: Graph theoretical analysis of structural and functional systems. Nature Rev Neurosci 10: 186–198. doi:10.1038/nrn2575.

Cao, H, Plichta, MM, Schffafer, A, Haddad, L, Grimm, O, Schneider, M, Esslinger, C, Kirsch, P, Meyer Lindenberg, A, Tost, H (2014) Test-retest reliability of fMRI-based graph theoretical properties during working memory, emotion processing, and resting state. Neuroimage 84: 888–900. doi:10.1016/j.neuroimage.2013.09.013.

Cicchetti, DV(1994) Guidelines, criteria, and rules of thumb for evaluating normed and standardized assessment instruments in psychology. Psychol Assess 3: 284–290. doi:10.1037/1040-3590.6.4.284.

Cocchi, L, Sale, MV, Lord, A, Zalesky, A, Breakspear, M, Mattingley, JB (2015) Dissociable effects of local inhibitory and excitatory theta-burst stimulation on large-scale brain dynamics. J Neurophysiol 113: 3375–3385. doi:10.1152/jn.00850.2014.

Colclough, GL, Woolrich, MW, Tewarie, PK, Brookes, MJ, Quinn, AJ, Smith, S (2016) How reliable are MEG resting-state connectivity metrics? Neuroimage 138: 284–293. doi:10.1016/j.neuroimage.2016.05.070.

Davis, SW, Kragel, JE, Madden, DJ, Cabeza, R (2012) The architecture of cross-hemispheric communication in the aging brain: Linking behavior to functional and structural connectivity. Cereb Cortex 22: 232–242. doi:10.1093/cercor/bhr123.

Deco, G, Jirsa, VK, McIntosh, AR (2011) Emerging concepts for the dynamical organization of resting-state activity in the brain. Nature Rev Neurosci 12: 43–56. doi:10.1038/nrn2961.

Delorme, A, Makeig, S (2004) EEGLAB: An open source toolbox for analysis of single-trial EEG dynamics including independent component analysis. J Neurosci Methods 134: 9–21. doi:10.1016/j.jneumeth.2003.10.009.

Desikan, RS, Sffegonne, F, Fischl, B, Quinn, BT, Dickerson, BC, Blacker, D, Buckner, RL, Dale, AM, Maguire, RP, Hyman, BT, Albert, MS, Killiany, R. J (2006) An automated labelling system for subdividing the human cerebral cortex on MRI scans into gyral based regions of interest. Neuroimage 31, 968–980. doi:10.1016/j.neuroimage.2006.01.021.

Deuker, L, Bullmore, ET, Smith, M, Christensen, S, Nathan, PJ, Rockstroh, B, Bassett, D.S. (2009) Reproducibility of graph metrics of human brain functional networks. Neuroimage 47: 1460–1468. doi:10.1016/j.neuroimage.2009.05.035.

Dosenbach, N. U. F, Nardos, B, Cohen, A. L, Fair, D. A, Power, J. D, Church, J. A, Nelson,S. M, Wig, G. S, Vogel, A. C, Lessov-Schlaggar, C. N, Barnes, K. A, Dubis, J. W, Feczko, E, Coalson, R. S, Pruett, JR, Barch, DM, Petersen, SE, Schlaggar, B. L (2010) Prediction of individual brain maturity using fMRI. Science 329: 1358–1361. doi:10.1126/science.1194144.

Fonov, V, Evans, AC, Botteron, K, Almli, CR, McKinstry, RC, Collins, DL (2011) Unbiased average age appropriate atlases for pediatric studies. Neuroimage 54: 313–327. doi:10.1016/j.neuroimage.2010.07.033.

Fornito, A, Zalesky, A, Bassett, DS, Meunier, D, Ellison-Wright, I, Yucel, M, Wood, SJ, Shaw, K, O’Connor, J, Nertney, D, Mowry, BJ, Pantelis, C, Bullmore, ET (2011) Genetic influences on cost-efficient organization of human cortical functional networks. J Neurosci 31: 3261–3270. doi:10.1523/JNEUROSCI.4858-10.2011.

Freeman, LC (1977) A Set of Measures of Centrality Based on Betweenness indicat. Sociometry, 4: 35–41.

Fuchs, M, Wagner, M, Kastner, J (2001) Boundary element method volume conductor models for EEG source reconstruction. Clin Neurophysiol 11: 1400{1407. doi:10.1016/S1388-2457(01)00589-2.

Geselowitz, D.B (1967) On bioelectric potentials in an inhomogeneous volume conductor. Biophys J 7: 1–11. doi:10.1016/S0006-3495(67)86571-8.

Gramfort, A, Papadopoulo, T, Olivi, E, Clerc, M (2010) OpenMEEG: opensource software for quasistatic bioelectromagnetics. Biomed Eng Online 9, 45. doi:10.1186/1475-925X-9-45.

Hagmann, P, Cammoun, L, Gigandet, X, Meuli, R, Honey, C.J, Van Wedeen, J, Sporns, O (2008) Mapping the structural core of human cerebral cortex. PLoS Biol 6: 1479–1493. doi:10.1371/journal.pbio.0060159.

Hardmeier, M, Hatz, F, Bousleiman, H, Schindler, C, Stam, CJ, Fuhr, P (2014) Reproducibility of functional connectivity and graph measures based on the phase lag index (PLI) and weighted phase lag index (wPLI) derived from high resolution EEG. PLoS ONE 9. doi:10.1371/journal.pone.0108648.

Horwitz, B (2003) The elusive concept of brain connectivity. Neuroimage 19, 466–470. doi:10.1016/S1053- 8119(03)00112-5.

Huang, YZ, Edwards, MJ, Rounis, E, Bhatia, KP, Rothwell, JC (2005) Theta burst stimulation of the human motor cortex. Neuron 45: 201–206. doi:10.1016/j.neuron.2004.12.033.

Jin, S. H, Seol, J, Kim, JS, Chung, CK (2011) How reliable are the functional connectivity networks of MEG in resting states? J Neurophysiol 106: 2888–2895. doi:10.1152/jn.00335.2011.

Kong, J, Gollub, RL, Webb, JM, Kong, JT, Vangel, MG, Kwong, K (2007) Test-retest study of fMRI signal change evoked by electroacupuncture stimulation. Neuroimage 34: 1171–1181. doi:10.1016/j.neuroimage.2006.10.019.

Kuntzelman, K, Miskovic, V (2017) Reliability of graph metrics derived from resting-state human EEG. Psychophysiology 54: 51–61. doi:10.1111/psyp.12600.

Langer, N, Pedroni, A, Jancke, L (2013) A next-generation approach to the characterization of a non-model plant transcriptome. PLoS ONE 8: 1–9. doi:10.1371/Citation.

Latora, V, Marchiori, M (2001) Efficient behavior of small-world networks. PRL 87: 1–4. doi:10.1103/PhysRevLett.87.198701.

Lord, A, Horn, D, Breakspear, M, Walter, M (2012) Changes in community structure of resting statefunctional connectivity in unipolar depression. PLoS ONE 7. doi:10.1371/journal.pone.0041282.

Lynall, M. E, Bassett, DS, Kerwin, R, McKenna, PJ, Kitzbichler, M, Muller, U, Bullmore, E(2010) Functional Connectivity and Brain Networks in Schizophrenia. J Neurosci 30: 9477–9487. doi:10.1523/JNEUROSCI.0333-10.2010.

Mahjoory, K, Nikulin, VV, Botrel, L, Linkenkaer-Hansen, K, Fato, MM, Haufe, S (2017). Consistency of EEG source localization and connectivity estimates. Neuroimage 152: 590–601. doi:10.1016/j.neuroimage.2017.02.076.

Newman, MEJ (2006) Modularity and community structure in networks. PNAS 103: 8577–8582. doi:10.1073/pnas.0601602103.

Nolte, G, Ziehe, A, Nikulin, VV, Schlogl, A, Kramer, N, Brismar, T, Muller, KR (2008) Robustly\estimating the ow direction of information in complex physical systems. PRL 100: 1–4. doi:10.1103/PhysRevLett.100.234101.

Nyffeler, T, Cazzoli, D, Hess, CW, Muri, RM (2009) One session of repeated parietal theta burst stimulation trains induces long-lasting improvement of visual neglect. Stroke 40: 2791–2796. doi:10.1161/STROKEAHA.109.552323.

Nyffeler, T, Wurtz, P, Luscher, H. R, Hess, CW, Senn, W, Pugshaupt, T, Von Wartburg, R, Luthi, M, Muri, RM (2006) Extending lifetime of plastic changes in the human brain. EJN 24: 2961–2966. doi:10.1111/j.1460-9568.2006.05154.x.

Oostenveld, R, Fries, P Maris, E, Schoffelen, JM (2011) FieldTrip: Open source software for advanced analysis of MEG, EEG, and invasive electrophysiological data. Comput. Intell. Neurosci 2011. doi:10.1155/2011/156869. arXiv:156869.

Pedersen, M, Omidvarnia, AH, Walz, JM, Jackson, GD (2015) Increased segregation of brain networks in focal epilepsy: An fMRI graph theory finding. Neuroimage: Clinical 8: 536–542. doi:10.1016/j.nicl.2015.05.009.

Rubinov, M, Knock, SA, Stam, CJ, Micheloyannis, S, Harris, AWF, Williams, LM, Breakspear, M (2009) Small-world properties of nonlinear brain activity inschizophrenia. Hum Brain Mapp 30: 403–416. doi:10.1002/hbm.20517.

Rubinov, M, Sporns, O (2010) Complex network measures of brain connectivity: Uses and interpretations. Neuroimage 52, 1059–1069. doi:10.1016/j.neuroimage.2009.10.003.

Rubinov, M, Sporns, O (2011) Weight-conserving characterization of complex functional brain networks. NeuroImage 56: 2068–2079. doi:10.1016/j.neuroimage.2011.03.069.

Schabrun, SM, Burns, E, Hodges, PW (2015). New insight into the time-course of motor and sensory system changes in pain. PLoS ONE 10: 1–14. doi:10.1371/journal.pone.0142857.

Schoffelen, JM, Oostenveld, R, Fries, P (2008). Imaging the human motor system’s beta-band synchronization during isometric contraction. Neuroimage 41: 437–447. doi:10.1016/j.neuroimage.2008.01.045.

Shrout, PE, Fleiss, JL (1979). Intraclass correlations: Uses in assessing rater reliability. Psychological Bulletin 86: 420–428. doi:10.1037/0033-2909.86.2.420.

Smith, SM, Nichols, TE, Vidaurre, D, Winkler, AM, Behrens, TE. J, Glasser, MF, Ugurbil, K, Barch, DM, Van Essen, DC, Miller, KL (2015). A positive-negative mode of population covariation links brain connectivity, demographics and behavior. Nat Neurosci 18: 1565–1567. doi:10.1038/nn.4125.

Sporns, O, Chialvo, DR, Kaiser, M, Hilgetag, CC (2004) Organization, development and function of complex brain networks. Trends Cogn Sci 8: 418–425. doi:10.1016/j.tics.2004.07.008.

Tadel, F, Baillet, S, Mosher, JC, Pantazis, D, Leahy, RM (2011) Brainstorm: A user-friendly application for MEG/EEG analysis. Comput Intell Neurosci 2011. doi:10.1155/2011/879716.

Tian, L, Wang, J, Yan, C, He, Y (2011) Hemisphere- and gender-related differences in small-world brain networks: A resting-state functional MRI study. Neuroimage 54: 191–202. doi:10.1016/j.neuroimage.2010.07.066.

Torta, DME, Van Den Broeke, EN, Filbrich, L, Jacob, B, Lambert, J, Mouraux, A (2017) Intense pain influences the cortical processing of visual stimuli projected onto the sensitized skin. Pain 158: 691–697. doi:10.1097/j.pain.0000000000000816.

Wang, L, Li, Y, Metzak, P, He, Y, Woodward, TS (2010) Age-related changes in topological patterns of large-scale brain functional networks during memory encoding and recognition. Neuroimage 50: 862–872. doi:10.1016/j.neuroimage.2010.01.044.

Watts, DJ, Strogatz, S. H (1998) Collectivedynamics of ‘small-world’ networks. Nature 393: 440–442. doi:10.1038/30918.

